# Demogenetic simulations reveal fragmenting effects of climate change on insular lizard populations

**DOI:** 10.1101/173922

**Authors:** Stephen E. Rice, Rulon W. Clark

## Abstract

The extinction risk of insular species with sessile life histories is expected to increase as they may be unable to track habitat in response to global climate change. Demogenetic simulations can couple population demography and niche modeling to produce spatially-explicit genetic and demographic information for all simulated individuals and provide insight into the effects of climate change at demographic and population genetic levels. We used CDMETAPOP to simulate a population of island night lizards (*Xantusia riversiana*) on Santa Barbara Island to evaluate its sensitivity to climate change to the year 2100 across 8 scenarios based on 2 climate models, 2 emissions pathways, and 2 connectivity models. We found that 1) *X. riversiana* is sensitive to climate change with SDMs predicting a loss of suitable habitat of 93%-98% by 2038, 2) population genetic structure is expected to increase drastically to 0.209-0.673 from approximately 0.0346, and 3) estimated minimum abundance is expected to declined sharply over the 2007 to 2038 period and reached values of 0-1% of the 2007 population size in all scenarios by 2100. Climate change is expected to decrease census population size and result in extant habitat patches that are isolated from one another with very high levels of genetic divergence over short periods of time. These patterns may drive the Santa Barbara Island population to extinction under certain scenarios. Management plans should address methods to improve connectivity on the island and attempt to create refugial patches. Contingency plans, such as translocation, may be required to prevent population extirpation. This study highlights the utility of demogenetic simulations in evaluating population demographic and genetic patterns under climate change with suggestions on workflows for running simulations in a high-throughput manner.

## INTRODUCTION

Global climate change may lead to rapid changes in environmental conditions, thus presenting unique challenges to species persistence. Climate change is expected to strongly affect species with restricted ranges, specialized niches, limited dispersal ability, and small effective population sizes (Oliver and Morecroft 2014), conditions intrinsic to many insular species. Populations may respond to climate change by tracking habitat shifts, coping through phenotypic plasticity or adaptation, or failing these, may go extinct (Penuelas et al. 2013). The persistence of populations is thereby directly related to organisms’ abilities to track suitable habitat or cope with changing conditions. For insular species, climate change may shift suitable niche space outside of the dispersal threshold and may result in sharp population declines, increased selective pressures, or extinction.

Species sensitivity to climate change has been assess through multiple modeling approaches, which range from species distribution models (SDMs) to stochastic simulations linking population demography to SDM predictions (e.g. Fordham et al. 2012; Swab et al. 2015). While effective tools for assessing population viability, these approaches do not incorporate genetic population structure, diversity estimates, and do not include scenarios for adaptation. Recent advances in individual-based simulations have led to landscape demogenetic models (Landguth et al. 2017a) which allow for a greater range of questions and scenarios to be investigated that incorporate demographic and genetic data.

Studies which use demogenetic models to investigate the effects of environmental change have been primarily focused on aquatic systems (e.g. Landguth et al. 2014; Piou et al. 2015). The use of detailed demogenetic models for climate change assessments in terrestrial systems remains largely unexplored. The flexibility of this modeling approach allows the coupling of population demography with SDM predictions to investigate the effects of climate change on population viability while also providing data to describe population genetic patterns. In addition to their flexibility, demogenetic models can simulate genetic data equivalent to empirical approaches which provide benchmarks for model parameterization (e.g. Row et al. 2014).

Islands provide exemplary systems for evaluating the use of demogenetic simulations in assessments of terrestrial systems due to their natural isolation and limited spatial extent. These attributes allow fine-scale simulations of closed populations to determine the sensitivity of insular species to climate change while simultaneously evaluating the practical constraints of demogenetic simulations. Recent research on the island night lizard, *Xantusia riversiana*, has characterized genetic patterns across two-thirds of the species range, found correlations of landscape features with genetic structure, and characterized dispersal patterns (Rice and Clark 2016, 2017) which provide empirical benchmarks for the parameterization of demogenetic models. Previous ecological research identified precipitation as an important variable in reproduction and calculated habitat-specific carrying capacities (Fellers and Drost 1991), suggesting a coupling of demography and climatic suitability. The island night lizard is one of the few reptiles endemic to the California Channel Islands; however, climate change sensitivity has yet to be assessed due to an expectation of limited effect on these locally abundant populations (United States Fish and Wildlife Service (USFWS) 2014). The combination of closed and well-characterized populations with the need for climate change sensitivity analyses presents a compelling opportunity to apply demogenetic simulations to this insular system.

We constructed demogenetic models in the program CDMETAPOP (Landguth et al. 2017a) for island night lizards on Santa Barbara Island (SBI) to determine the effects of climate change on expected minimum abundances (EMA), quasi-extinction risks, and population genetic structure. We used stochastic simulations which included demographic and environmental stochasticity to assess population sensitivity to climate change. We modeled the effects of climate change through annual changes in patch carrying capacity and evaluated the sensitivity of extinction risk and population genetic patterns to variability in the effective distances between patches due to climate change. Finally, we evaluated the practicality of using demogenetic models to conduct these analyses and identified approaches to improve these models in terrestrial systems.

## METHODS

### Study Species

*Xantusia riversiana* is the only reptile endemic to three California Channel Islands: SBI, San Clemente Island (SCI), and San Nichols Island. Each island is a distinct evolutionary and management unit (USFWS 2014). Two subspecies were recognized by Smith (1946), with SBI and SCI populations grouped as *X. r. reticulata* and San Nichols Island recognized as *X. r. riversiana* based on morphology. In addition to morphology, the San Nichols Island population differs in habitat utilization, body size, and reproductive rates (Fellers et al. 1998). Island night lizards are habitat generalist on SBI and SCI with the greatest densities occurring in California boxthorn (*Lycium californicum*), prickly pear cactus (*Opuntia littoralis*), and rocky habitats (Fellers and Drost 1991; Mautz 1993). Population densities in prime habitat are estimated to be in excess of 3,200 individuals/ha with total population sizes on SCI estimated at 21.3 million and 17,600 on SBI (USFWS 2014). Individuals on SCI may live in excess of 23 years (Mautz 2015, pers. comm) with sexual maturity reached between 2 and 3 years (Fellers and Drost 1991; Goldberg and Bezy 1974). Reproduction is is influenced by precipitation patterns and hypothesized to occur biennially for mature females (Fellers and Drost 1991). Geographic distance and landscape features were correlated with genetic distance on SBI and SCI (Rice and Clark 2016). Landscape features included negative correlations with prime habitat and positive correlations with canyons, secondary roadways, and coastal cholla cactus (*Cylindropuntia prolifera*) on SCI and wooly seablite (*Suaeda taxifolia*), crystalline iceplant (*Mesembryanthemum crystallinum*), and barren ground on SBI. Estimates of dispersal distance range from an average displacement of 3m (Mautz 1993) to genetically inferred dispersal ranging from 14m on SBI to 41m on SCI (Rice and Clark 2017). Island night lizards were delisted from threatened status under the Endangered Species Act in 2014, contingent on post-delistment monitoring. Climate change sensitivity has not been assessed for the species but is anticipated to be of minimal impact (USFWS 2014).

### Habitat Suitability Models

We constructed SDMs in MAXENT (Phillips et al. 2006) to identify contemporary correlates of climatic niche with occurrences limited to SCI and SBI, due to shared ecological and life history patterns. We used occurrence data from capture data (Rice and Clark 2016) and GBIF (doi:10.15468/dl.rie3zo). We removed records without GPS coordinates, those that mapped outside of SBI and SCI, and those prior to 1959 to limit occurrence records to those which potentially overlap historic climate data. The occurrence data of Rice and Clark (2016) were spatially biased by distance to trails (SBI) and distance to roadways (SCI); therefore, we constructed bias maps using the distance of a cell to primary access route to assign probabilities of sampling a given cell. The probability of a given cell being sampled was the proportion of the cell’s distance bin among all raster cells for each respective island.

Climatic predictors were the first 19 bioclim variables generated from 800 m^2^ historical (1981-2010) PRISM precipitation and temperature data (PRISM 2012) using the R package DISMO (Hijmans et al. 2017). Terrain predictors of slope, aspect, terrain roughness, terrain ruggedness, and topographic position were constructed from a 10 m resolution National Elevation Dataset (United States Geological Survey 2017) and resampled to 100 m using the R package RASTER (Hijmans et al. 2016). Highly correlated predictors were removed through stepwise variance inflation factor (VIF) analysis in the R package USDM (Naimi 2015) with a threshold of 10. Cleaned occurrence data was used with retained predictors to construct SDMs using the autofeatures setting, duplicate occurrence records removed, and bootstrapped 1000 times (Franklin 2010).

Two global climate models were chosen for the projection of SDMs to future time periods. The two models, CanESM and Miroc, are considered predictive for the Basic Characterization Model, which informs many California climate assessments (Flint et al. 2013). The CanESM and Miroc climate models, at representative concentration pathways (RCPs) of 4.5 and 8.5, were downscaled to 100 m resolution by Dr. Alan Flint (USGS) for monthly temperature and precipitation variables. We constructed bioclim variables for projections to the years 2038, 2069, and 2100 by averaging monthly variables over the 30 years prior to each time point and using these averages to produce bioclim predictors. Terrain variables remained unchanged in all scenarios. The bootstrapped SDMs were projected to each time point and averaged to identify habitat suitability under climate change scenarios. We defined the contemporary habitat suitability map as the year 2007 for all downstream analyses and interpolations, which allowed consistent 30 year time spans for all endpoint projections.

### Demogenetic Simulations

We modeled the effects of climate change on population viability and genetic structure using the demogenetic modeling approach of Landguth et al. (2017) with the program CDMETAPOP. Demogenetic models included climate change as a process which modified the habitat suitability of each raster cell. The effect of climate change on population viability was investigated by linking patch carrying capacities to habitat suitability values. Additionally, we parameterized two resistance surface models to evaluate the effect of climate change on inter-patch connectivity. Two resistance surface models were evaluated, a static model based on geographic distance and a dynamic model based on habitat suitability. We linearly interpolated between each endpoint to produce annual SDMs ranging from 2007 through the year 2100 using RASTER. We modeled every 1 ha raster cell as a distinct patch with a constant relationship between carrying capacity and habitat suitability. The constant relationship between individuals and habitat suitability was determined by taking the census population size and dividing by the sum of contemporary habitat suitability. Populations on SBI were modeled with patch carrying capacities determined by habitat suitability multiplied by the constant of 117.76 individuals.

Demographic parameters were drawn from the ecological literature of *X. riversiania* (Fellers and Drost 1991; Mautz 1993) and *X. vigilis* (Zweifel and Lowe 1966). *Xantusia riversiana* exist in age structured populations; however, there is insufficient data to estimate the survival parameters needed to model demography. We used a 4-stage model (Table S1) of demography informed by survival estimates of a sister species, *X. vigilis*, which occur on the mainland. Both species display similar patterns of high juvenile survival (Mautz 1993) and reproductive potential (Goldberg and Bezy 1974). Fecundity values for *X. riversiania* from Fellers and Drost (1991) were used as parameters (Table S1) for reproductively active (4^th^ stage) females which were modeled as seasonally monogamous with strict biennial reproduction.

All models included environmental and demographic stochasticity. Environmental stochasticity was incorporated through standard deviations in patch carrying capacity. The standard deviation for each patch at each time step was determined by applying the proportion of the standard deviation to mean population density on San Clemente Island, 0.1415 (Mautz 1993). Demographic stochasticity was modeled through all time steps as variability in survival rates and fecundity. Survival rate variability was modeled through the standard deviation in survival rates for each age group as estimated from Table 4 in Zweifel and Lowe (1966). Variability in fecundity was modeled by assigning the offspring number for each female from a normal distribution around the mean with standard deviation (Fellers and Drost 1991).

We used the same cost distance matrices and dispersal formula parameters and thresholds for mating and movement for both sexes. The cost distance matrices for each time step consisted of the effective resistance distances between all patches as calculated by the program CIRCUITSCAPE (McRae and Beier 2007). The static model based on geographic distance was constructed with a raster map with all terrestrial cells assigned a neutral value of 1, which constrained dispersal paths to land and has been shown to approximate log-transformed geographic distance (Lee-Yaw et al. 2009). Dynamic resistance maps, which assessed changes in connectivity, were produced for each time step with resistance set as the inverse of the habitat suitability for each raster cell. We modeled movements with a negative exponential function (scale =1.0, shape =0.75) and a maximum effective resistance distance threshold of 0.555 for geographic distance models and 1.25 for connectivity models. These values were chosen as they resulted in equilibrated global Fst estimates similar to the observed empirical value of 0.0346 (Rice and Clark 2016) when modeled on the contemporary suitability map for 450 years. In addition to the 4 climate change scenarios, we modeled a “no change” scenario in which contemporary habitat suitability remained the same throughout the simulation.

We simulated 17,600 individuals with 20 microsatellite loci each with 15 alleles and no mutation. Simulations began with a panmictic population and equilibrated on contemporary conditions for 450 years to produce a 2007 estimate of global Fst approximate to the empirical data. Changes in carrying capacity and effective resistance distance began at the 2008 time step. Populations were fully sampled at the years 2038, 2069, and 2100 and global Fst calculated in the R package DIVERSITY (Keenan et al. 2013). Full census data was extracted for each timestep between 2007 and 2100 and was used to calculate proportional EMA, compared to 2007, and generate quasi-extinction graphs following McCarthy and Thompson (2001). All scenarios were replicated 100 times and ran on a high performance computational (HPC) cluster.

## RESULTS

### Habitat Suitability

Seven Bioclim variables and 3 terrain variables were retained by stepwise VIF (Table 1) resulting in a maximum correlation between Bio5 and Bio9 of 0.7706. Maximum temperature in the warmest month, Bio5, had the most useful information by itself and topographic position index (TPI) had the most information not present in other variables. The minimum observed habitat suitability value for any occupancy record was 0.1095, which was used as the threshold definition to visualize habitat patches (Supplemental Figures S1-S13) and differentiate suitable from unsuitable habitat patches. All climate scenarios displayed striking decreases in suitable habitat availability and distribution. These models predict 93.13%-99.81% reductions in suitable habitat on SBI and SCI from 2007 under all scenarios for the year 2038 (Figure 1), which correspond to a change from 7482 ha under current conditions to 514 - 14 ha under climate change. Suitable habitat expanded on a nearby island to the east of SBI and SCI, Santa Catalina, which is outside of the current and historic species distribution. The Miroc 8.5 model retained the most suitable habitat on SCI and SBI combined. Habitat suitability for all scenarios declined through the year 2069 and declined to 0 suitable hectares under both CanESM scenarios and the Miroc 4.5 scenario by the year 2100. The Miroc 8.5 scenario was the only scenario investigated which did not result in the complete elimination of suitable habitat.

**Table 1:**
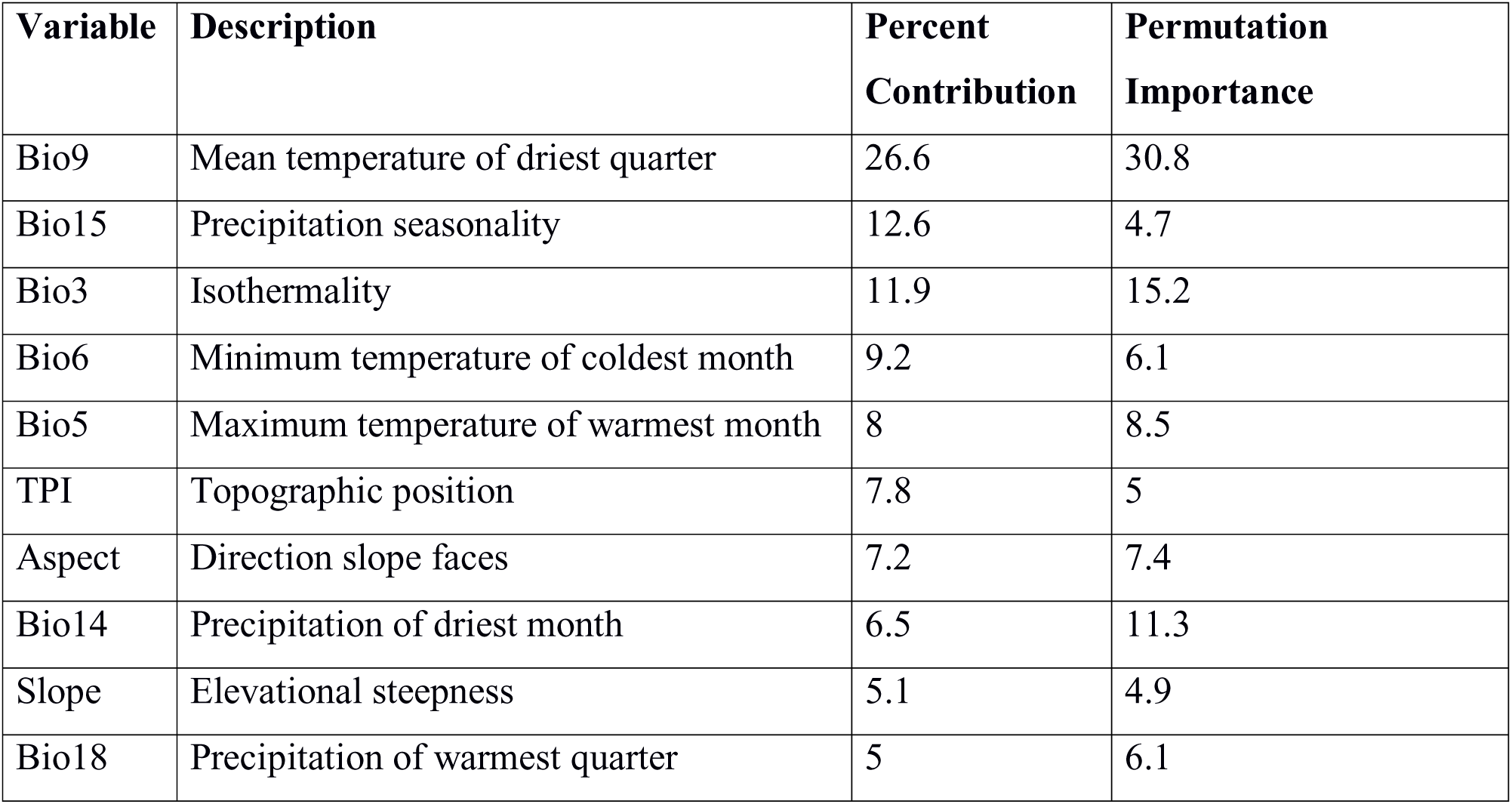
Environmental and topographic predictors. Predictors were retained for models of SBI and SCI based on stepwise VIF. MAXENT calculated percent contribution and permutation importance for each variable.

| Variable | Description | Percent Contribution | Permutation Importance |
| --- | --- | --- | --- |
| Bio9 | Mean temperature of driest quarter | 26.6 | 30.8 |
| Bio15 | Precipitation seasonality | 12.6 | 4.7 |
| Bio3 | Isothermality | 11.9 | 15.2 |
| Bio6 | Minimum temperature of coldest month | 9.2 | 6.1 |
| Bio5 | Maximum temperature of warmest month | 8 | 8.5 |
| TPI | Topographic position | 7.8 | 5 |
| Aspect | Direction slope faces | 7.2 | 7.4 |
| Bio14 | Precipitation of driest month | 6.5 | 11.3 |
| Slope | Elevational steepness | 5.1 | 4.9 |
| Bio18 | Precipitation of warmest quarter | 5 | 6.1 |

**Figure 1:**
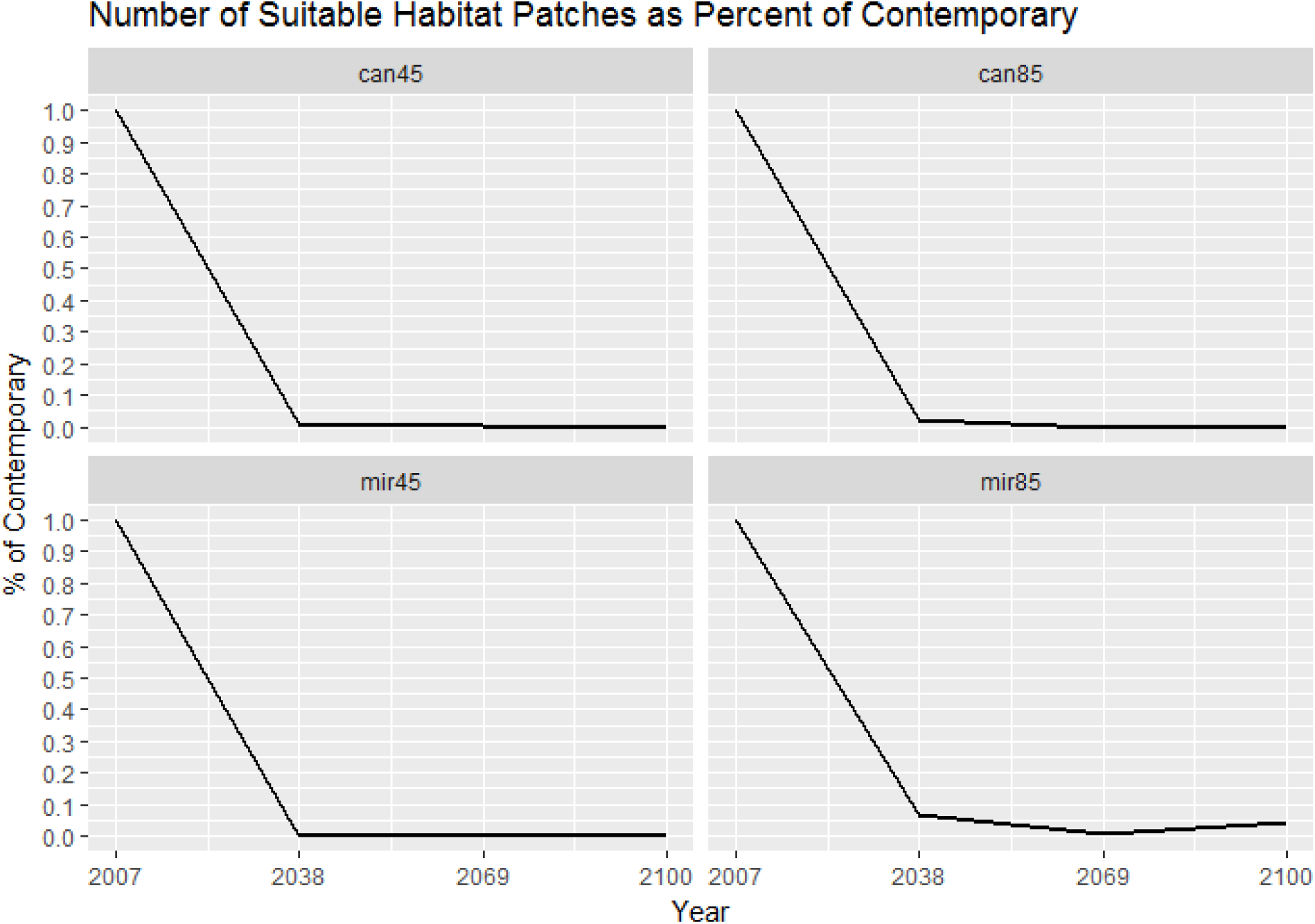
Percentage of Contemporary Suitable Habitat Patches Remaining on both SBI and SCI. Habitat patches were defined based on an observed occupancy value of 0.1095 with the % of contemporary values representing percentages 0 to 1.

### Demogenetic Simulations

All models equilibrated at global Fst values (Figure 2) similar to the empirical data of Rice and Clark (2016). The distance only models revealed sharp increases in global Fst across all climate scenarios. Mean global Fst was approximately 0.043 at the 2007 time step in equilibrated models and increased to a range of 0.209-0.330 by 2100 depending on the model (Figure 2, top). The CanESM 8.5 model could not be assessed at the year 2100 due to population extinction throughout the simulations. EMA revealed sharp population declines (Figures 3-5, top) at all time points with the Miroc 8.5 projection being the most optimistic with 1.76% of the 2007 population remaining by 2100 (Table 2). The CanESM 8.5 model resulted in complete extinction and the remaining models ended the century with ≤ 0.15% of the 2007 population remaining. Quasi-extinction graphs (Figures 3-5, top) also show increased extinction risks at 2038 with sharp increases in extinction risk through the end of the century. Connectivity models revealed the same trends and model rankings in global Fst as distance models; however values were almost twice as great (Figure 2, bottom). Mean global Fst values for connectivity models ranged from 0.5318-0.6733 by 2100 with CanESM 8.5 absent due to extinction. EMA and quasi-extinction analyses for connectivity models returned the same trends as distance models at similar values with the CanESM 8.5 model again resulting in extinction (Table 2, Figures 3-5, bottom).

**Figure 2:**
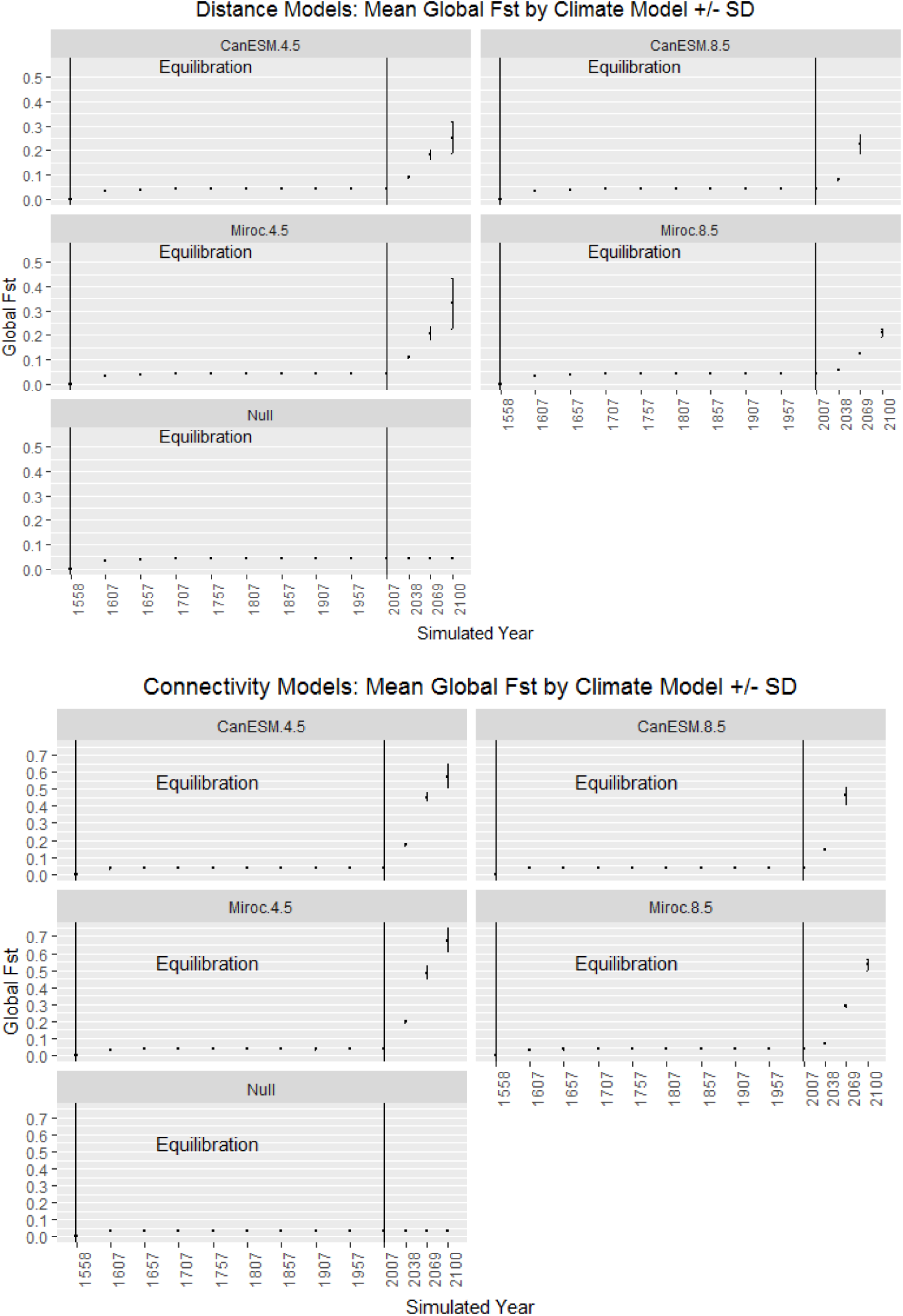
Mean global Fst with standard deviations. Distance-based models (top) and connectivity models (bottom) listed with climate model identification with RCP is given in the title. The Y-axis is global Fst averaged over all simulations for each model with error bars indicating standard deviation of the mean. Vertical lines denote the range considered for model equilibrium. Mean Fst is denoted with +/- standard deviations plotted from point data from sampled time points (X-axis).

**Figure 3:**
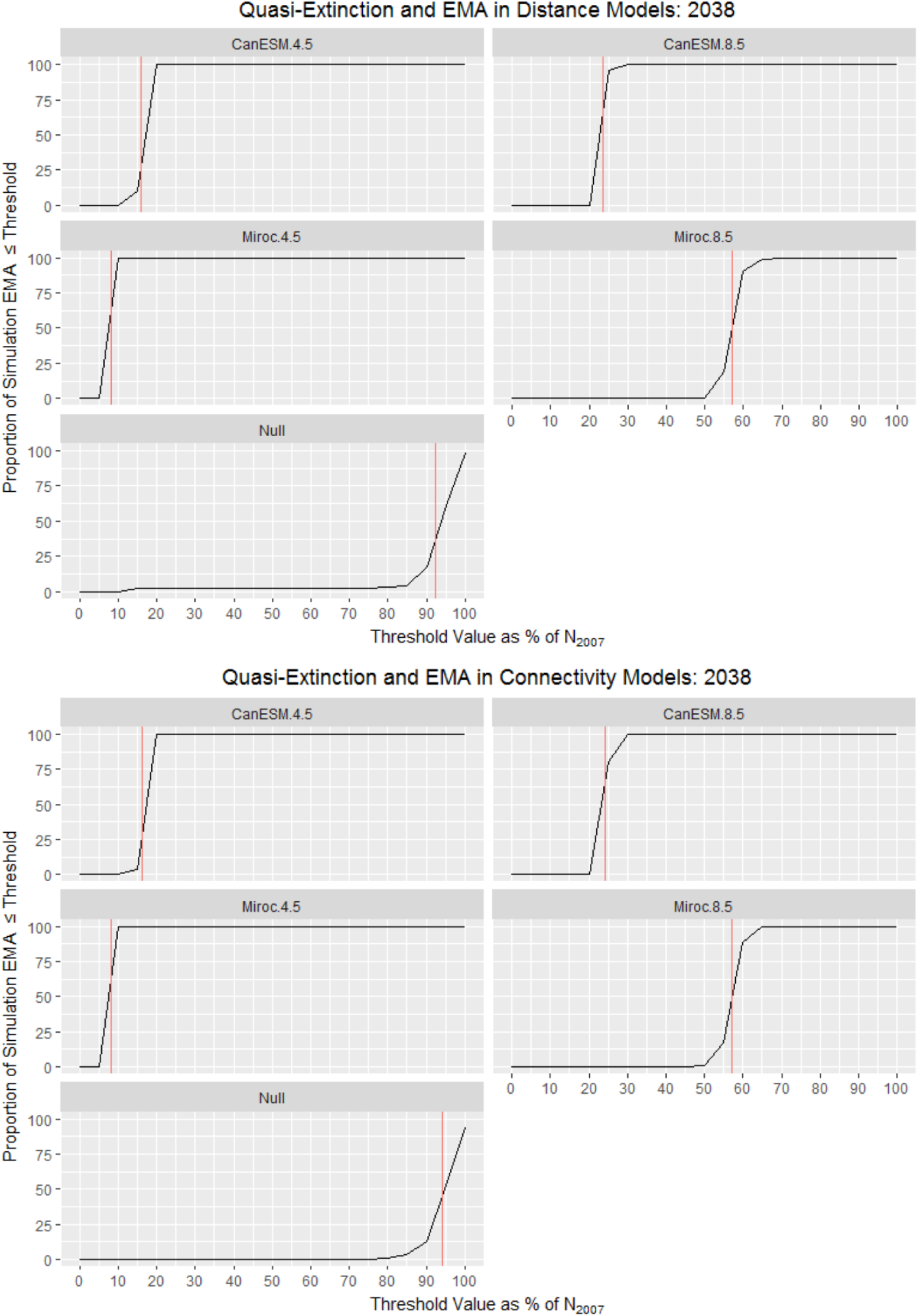
Quasi-extinction for distance and connectivity models 2007-2038. Distance-based models (top) and connectivity models (bottom) listed with climate model identification with RCP is given in the title. The Y-axis represents risk, defined as the proportion of models with a minimum size, represented as the proportion of the 2007 census population size remaining, less than or equal to the threshold value (X-axis). The red vertical line represents the proportional EMA averaged across all runs for a given model.

**Figure 4:**
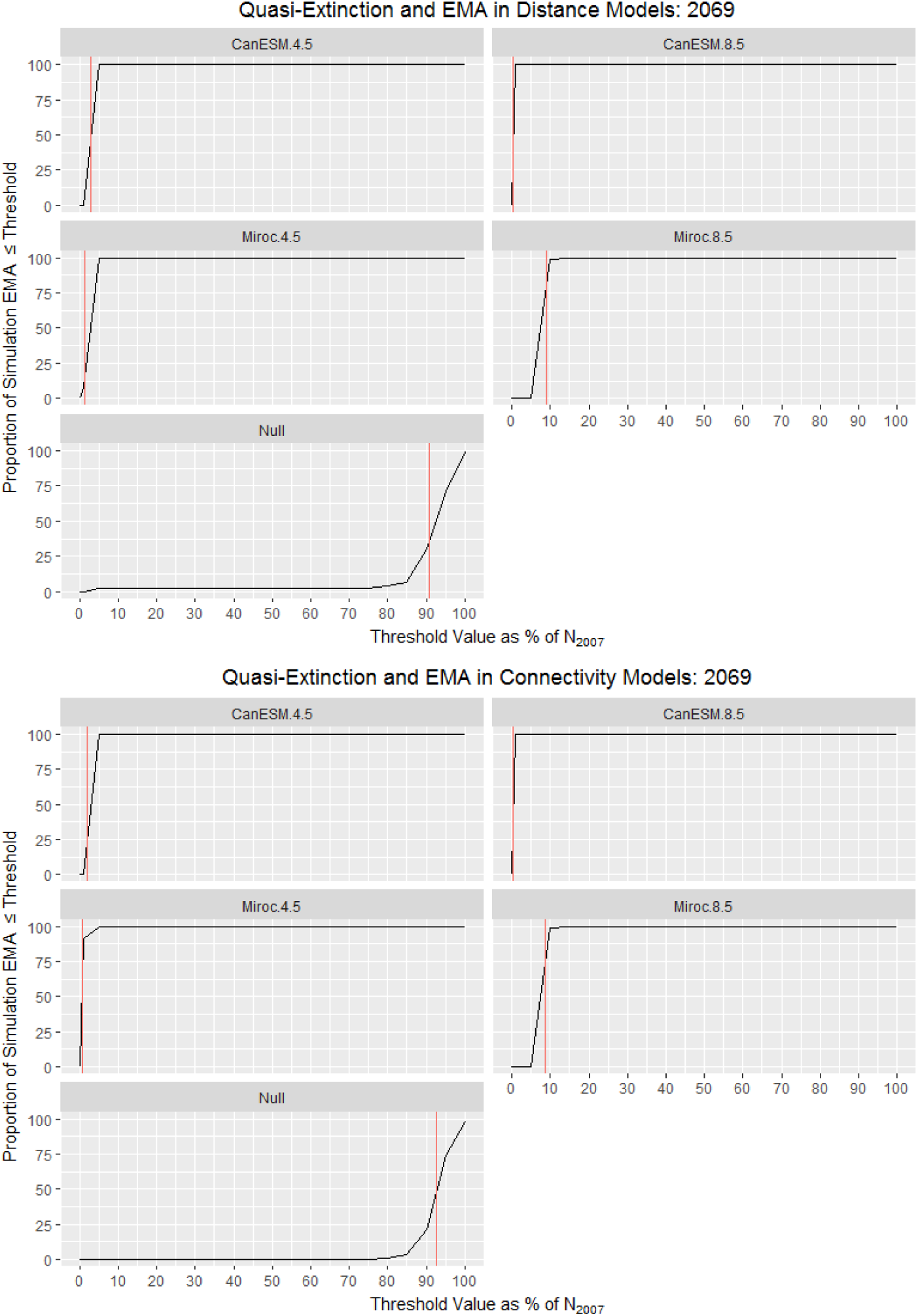
Quasi-extinction for connectivity models 2007-2069. Distance-based models (top) and connectivity models (bottom) listed with climate model identification with RCP is given in the title. The Y-axis represents risk, defined as the proportion of models with a minimum size, represented as the proportion of the 2007 census population size remaining, less than or equal to the threshold value (X-axis). The red vertical line represents the proportional EMA averaged across all runs for a given model.

**Figure 5:**
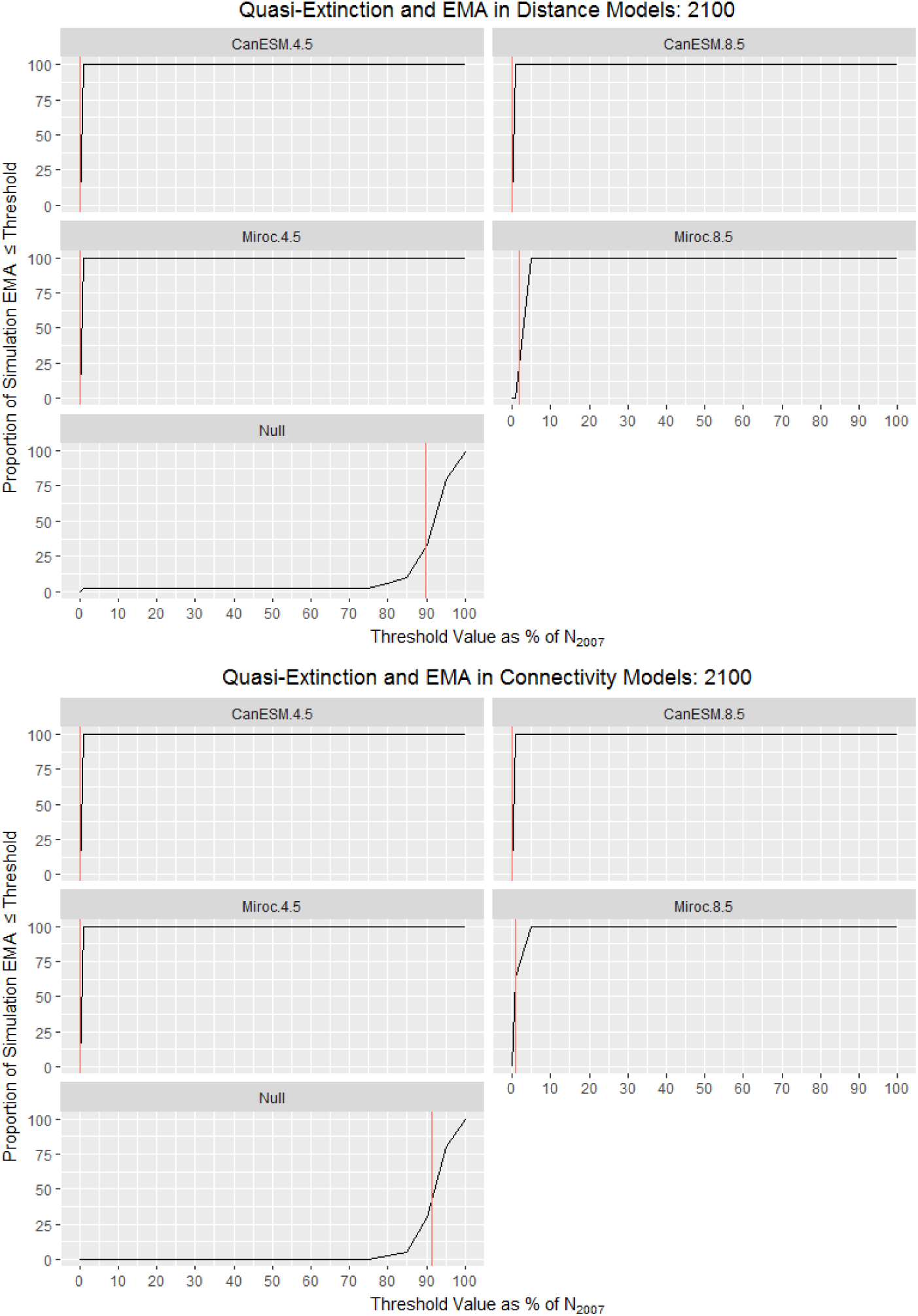
Quasi-extinction for connectivity models 2007-2100. Distance-based models (top) and connectivity models (bottom) listed with climate model identification with RCP is given in the title. The Y-axis represents risk, defined as the proportion of models with a minimum size, represented as the proportion of the 2007 census population size remaining, less than or equal to the threshold value (X-axis). The red vertical line represents the proportional EMA averaged across all runs for a given model.

**Table 2:** Proportional EMA by year, climate change scenario, and model. Simulation averaged proportional EMA between the observed minimum abundance between the contemporary model (2007) and the endpoint, over 100 stochastic simulations. Climate models were CanESM and Miroc with 2 RCPS, 4.5 and 8.5, and a null model of stable contemporary conditions. The distance model refers to the stable geographic equivalent resistance model and connective model refers to variable effective distances between patches.

|  | Distance Model |  |  | Connectivity Model |  |  |
| --- | --- | --- | --- | --- | --- | --- |
| Climate Model | 2038 | 2069 | 2100 | 2038 | 2069 | 2100 |
| CanESM 4.5 | 15.790 | 2.819 | 0.150 | 16.181 | 1.961 | 0.082 |
| CanESM 8.5 | 23.403 | 0.379 | 0.000 | 24.175 | 0.299 | 0.000 |
| Miroc 4.5 | 8.065 | 1.354 | 0.094 | 8.269 | 0.747 | 0.044 |
| Miroc 8.5 | 57.000 | 8.828 | 1.760 | 57.139 | 8.696 | 0.924 |
| No Change | 92.920 | 90.688 | 89.840 | 94.056 | 92.747 | 91.547 |

## DISCUSSION

SDMs can offer insight into the predicted shifts in habitat suitability for a species under climate change projections assuming that contemporary correlations hold (Franklin 2010). When combined with demogenetic simulations, these models allow for a spatially-explicit investigation into the effects of climate change on population demographic and genetic patterns. The current monitoring framework for the island night lizard suggests that climate change is not a concern for this species (USFWS 2014); however, our results indicate this species may be heavily impacted by climate change with reductions in suitable habitat and steep declines in population abundance. Under all climate change scenarios, SDMs predict a 93.13%-99.81% loss of suitable habitat on SBI and SCI. We investigated the sensitivity of *X. riverisiana* to climate change on SBI following a coupled niche-population model framework to determine the effects of climate change on population abundance and population genetic structure. We found global Fst values increased from the 2007 time point by factors of 4.76-7.55 for models with only geographic distance and 14.23-18.31 for models where climate change modified the intervening matrix. EMA revealed stark decreases in abundances across all climate models beginning in the year 2038 and extending until the end of the century. EMA at the year 2100 reached extinction for the CanESM 8.5 scenario and a maximum of 1.76% of the 2007 population under the Miroc 8.5 model with a distance only dispersal scenario. EMA values for other climate models were <1% by 2100 (Table 2).

### Habitat Suitability

The SDMs for *X. riversiana* represent the first efforts to characterize changes in habitat suitability for reptiles on the California Channel Islands. These models were correlative models built with abiotic factors and thus do not include vegetation or soil characteristics which may contribute towards the true suitability of habitat to support island night lizard populations. MAXENT is considered one of the most accurate SDM methods but may lead to more pessimistic projections than other methods (Conlisk et al. 2013). Even with these limitations, the contemporary SDM offered key insights into climatic factors which correlate with *X. riversiana* distributions on SBI and SCI. Precipitation is positively correlated with reproduction and recruitment on island and mainland *Xantusia* species (Fellers and Drost 1991; Zweifel and Lowe 1966). Precipitation was incorporated in this SDM through the bioclim suite of variables which include interactions between temperature and precipitation. The largest contributing variable to these models was Bio9, the mean temperature in the driest quarter, which corresponded to summer, June through August, on the California Channel Islands and overlaps with the gestational period of *X. riversiana* (Goldberg and Bezy 1974). The climatic niche and tolerances of *X. riversiana* are poorly understood, including the effects of temperature and water availability on successful gestation, and this variable may represent physiological constraints associated with gestation and realized climatic niche. Further experimental research is needed to examine the hypothesis that reproductive physiology and environmental constraints on gestation limit this species climatic niche.

SDMs projected under the Miroc and CanESM climate models revealed similar trends across both RCPs. The greatest decline in suitable habitat occurred between the contemporary model and 2038. The steepness of this decline, a loss of approximately 93% - 99% of contemporary habitat, is jarring given the longevity of the species and recent delistment. While these values may be pessimistic, they offer a clear indication that climate change is projected to have much larger effects on insular species within the California Floristic Province than included in current management plans. Based on the sensitivity of this highly abundant insular reptile, we recommend that species management and conservation efforts on the California Channel Islands incorporate habitat suitability forecasts, as habitat tracking may be an inviable option for strictly terrestrial species which may require translocation and assisted colonization to prevent extinction under climate change.

### Demogenetic Simulations

Individual-based spatially explicit demogenetic models are a recent tool which may give insight into processes generating population genetic patterns and population demographic trajectories. Previous research with CDMETAPOP is limited due to the recent publication of the method (Landguth et al. 2017a) but has included evaluation of blister rust resistance scenarios in whitebark pine (Landguth et al. 2017b) and invasive species management (Landguth et al. 2017a). However, CDMETAPOP is built modularly to draw on previous individual-based spatially-explicit models, such as CDPOP (Landguth and Cushman 2010), which have been utilized to explore topics ranging from climate change sensitivity in Lynx (Row et al. 2014) to experimental design (Rico 2017). Demogenetic simulations conducted with CDMETAPOP may be parameterized to yield coupled niche-population models with genetic data which can enhance research into the effects of connectivity, population structure, or adaptation under changing climate. Demogenetic simulations may improve coupled niche population modeling efforts by providing greater insight into the effects of climate change but the tradeoffs of this approach include increased computational time, computational load, and can be labor intensive to parameterize and analyze.

We approached demogenetic simulations from the framework of coupled niche population models by linking habitat suitability values with patch carrying capacity and varying that carrying capacity at each time step. Additionally, we explored the effect of modeling effective resistance between patches as a function of annual habitat suitability and compared this approach to a more traditional approach assuming geographic distance alone. We found demogenetic simulation to be a computationally expensive tool that provided insight into population viability and genetic structure under multiple climate change and effective distance scenarios. The scale of this study was only possible through the use of a HPC cluster to increase model throughput during parameterization and simulation, and would have been intractable on a standard workstation due to processing time and the storage space required for simulations and analyses (>1 TB for this study). The ability to run CDMETAPOP across multiple cores of a computer cluster and deploy basic R scripts to adjust model parameters and analyze simulated data allowed model parameterization and simulations to occur in a high-throughput manner which was only limited by computer usage quotas. Demogenetic models constructed for SBI were 17,600 individuals spread across 247 1 ha patches for a time period 542 years. A single iteration could complete on a single processor with 2GB of memory in 8-12 hours. This partitioning of computational load across multiple cores allowed an increase in the number of model runs, but still only a fraction of the number used in traditional coupled niche population models (e.g. Conlisk et al. 2013, Swab et al. 2015). However, the number was similar to other demogenetic approaches (e.g. Landguth et al. 2014; Piou et al. 2015). The parameterization and execution of simulations on a computer cluster is recommended to ensure robust examination of parameter space, and will be essential if modeling changes in genetic patterns from an assumed equilibrium in species of high density and continuous distribution.

Failure to validate equilibrium conditions for global Fst, and presumably other genetic metrics, when parameterizing models may lead to spurious and incorrect inferences. The current configuration of CDMETAPOP prevents parallelization and we found that the computational load becomes intractable for millions of individuals. Our attempts to parameterize models of SCI revealed that even at 10% of the census population size (2.15 million) two computer cores given 15 GB of RAM on a computer cluster were unable to generate the initial population in a 24 hour period. Furthermore, we found that efforts to parameterize models based on reducing the modeled population to the effective population size at each collection site failed to reach equilibrium of global Fst values even when dispersal was set to the maximum threshold. While we could achieve values equivalent to empirical values, these values were not stable and continued to increase even as effective distance and carrying capacity remained stationary. Failure to validate the assumed pattern or compare would have led to a serious inference error on the magnitude of change expected under climate change scenarios.. In addition to validating equilibrium conditions when examining questions of genetic structure, researchers may be forced to examine various simulated population sizes or spatial extents based on computational constraints.

### Implications for Conservation

The current distribution of *X. riversiana* will likely constrain its ability to cope with climate change through habitat tracking due to insular populations and dispersal distances under 50 m on both SBI and SCI. The projected size of habitat available to island night lizards based on SDM predictions is concerning due to the sharp declines in suitable habitat by 2038 in all scenarios and continued decline through the end of the century. Conservation efforts on both islands should focus on the creation of refugia/management areas in regions predicted to be the most suitable across climate predictions in an effort to ameliorate the anticipated loss of suitable habitat over the next 2 decades. Demogenetic simulations on SBI revealed populations are sensitive to climate change across all climate change scenarios when climate influences carrying capacities. We found that global Fst as a metric of intra-island isolation of patches were more sensitive to the resistance of the intervening matrix whereas demographic patterns represented by EMA were not. Rice and Clark (2016) found landscape correlates with genetic distance on SCI and SBI in addition to geographic distance. Simulation levels of global Fst suggest that even in scenarios of stable geographic distance, isolation of populations poses a major concern as global Fst values corresponded to significant population structure (Figure 2). These findings suggest that populations may be threatened by climate change through habitat loss, habitat degradation, and isolation of suitable patches. Increased isolation of patches even within a small island may further increase the extinction risk of the species through inbreeding depression or Allee effects. Based on SDM and simulation predictions, management intervention may be required to prevent extinction of *X. riversiana* on SBI as early as 2038. While we were unable to model SCI in demogenetic simulations, we expect similar patterns to hold given the shared life history traits and prime habitat requirements. The census population size is greater on SCI than SBI, thus the risk of quasi-extinction is likely reduced but sharp declines in population abundance are anticipated based on the loss of suitable habitat indicated by SDMs (Figures S1-S13).

Recommended management interventions include increasing prime habitat of California boxthorn and prickly pear cactus in areas of greater projected habitat suitability and between populations identified by Rice and Clark (2016) to serve as potential refugia and improved connectivity among populations. As a generalist species, the simulated responses of *X. riversiana* on SBI may be indicative of increased threats to plant communities or endemic deer mouse populations from climate change and a lack of viable habitat tracking. This study is specific to island night lizards, but highlights a need for further study among insular species for which climate change sensitivity analyses are lacking.

### Conclusions

Based on the results of demogenetic modeling, island night lizards are at high risk from climate change over the next several decades. The demographic effects of climate change over a range of emission and model scenarios are predicted to cause sharp population declines through decreased carrying capacities and sharp increases in population structure even when only geographic distance between patches is considered. We recommend that the National Park Service, the managing entity of SBI, and other agencies continue to monitor habitat and population sizes and key climatic variables to determine if populations begin to decline as predicted by our models. Additional factors may add to persistence of *X. riversiana* populations under climate change, such as responses of vegetation or soil characteristics necessary to maintain thermoregulation or unidentified plasticity within the species that could buffer against declines. These modeling results are also subject to the limitations of the method, chiefly that we assume the niche of island night lizards is well described by environmental variables, that contemporary relationships hold in future scenarios, and that population demography is monotonically coupled to habitat suitability. While these assumptions are likely violated, further research is needed into the sensitivity of model predictions to changes in population size, SDM construction, and parameters associated with vital rates. Additional research with demogenetic simulations within this system should address hypotheses related to adaptive responses to heat and drought tolerance, which can be implemented within the demogenetic models.

Demogenetic models are a valuable tool for climate change sensitivity analyses, but there usage is constrained to populations consisting of tens of thousands. Parameterization of models to yield genetic patterns which approximate empirical conditions under equilibrium settings can highlight model sensitivities to the spacing and densities of populations while providing benchmarks for model parameterization. We found stochastic simulations were informative to evaluate the sensitivity of an island population to environmental and demographic stochasticity as well as the permeability of the intervening matrix and provide insight into the genetic and demographic trajectories this species may face during the remainder of the century. While 100 stochastic simulations is below the traditional population viability analyses, it is equivalent to previous work with demogenetic simulations. Leveraging HPC clusters should alleviate some of the computational constraints for more robust simulation sizes. We recommend researchers interested in demogenetic simulations pursue HPC solutions to allow for increased simulations and robust model inferences. Secondary constraints are the computational load of analyzing simulated data and will vary with the research being conducted. Estimation of extinction risk and temporal trends in population size are easily extracted form summary tables within simulations; however metrics of population structure, such as global Fst, require more effort and considerably more time to extract. As demogenetic simulations grow in usage, dedicated workflows will need to emerge to reduce computational loads and provide an implementation to conduct thorough sensitivity analyses on the effect of model parameters.

## ACKNOWLEDGEMENTS

Support for this research came from United States Department of Defense (Award Number W9126G-12-2-0060) and the Southern California Research and Learning Center (Award Number S18309). We would like to thank the National Park Service for access to SBI, SCI Naval Base for access to SCI and our field assistants. In addition we would like to thank Drs. A. Bohonak (SDSU), H. Regan (UCR), K. Anderson (UCR), and J. Gatesy (UCR) for comments received during manuscript preparation.

---

**Table S1:** Demographic model

| **Age Class** | **Mortality**<br>**(SD)** | **Male**<br>**Maturity** | **Female**<br>**Maturity** | **Fecundity**<br>**(SD)** |
| --- | --- | --- | --- | --- |
| 0 | 11.7 (13.45) | 0 | 0 | 0 |
| 1 | 16.3 (16.05) | 0 | 0 | 0 |
| 2 | 19.5 (21.15) | 0.5 | 0 | 0 |
| 3 | 15.6 (18.63) | 1.0 | 1.0 | 3.9 (1.15) |

**Figures: S1-S13.**
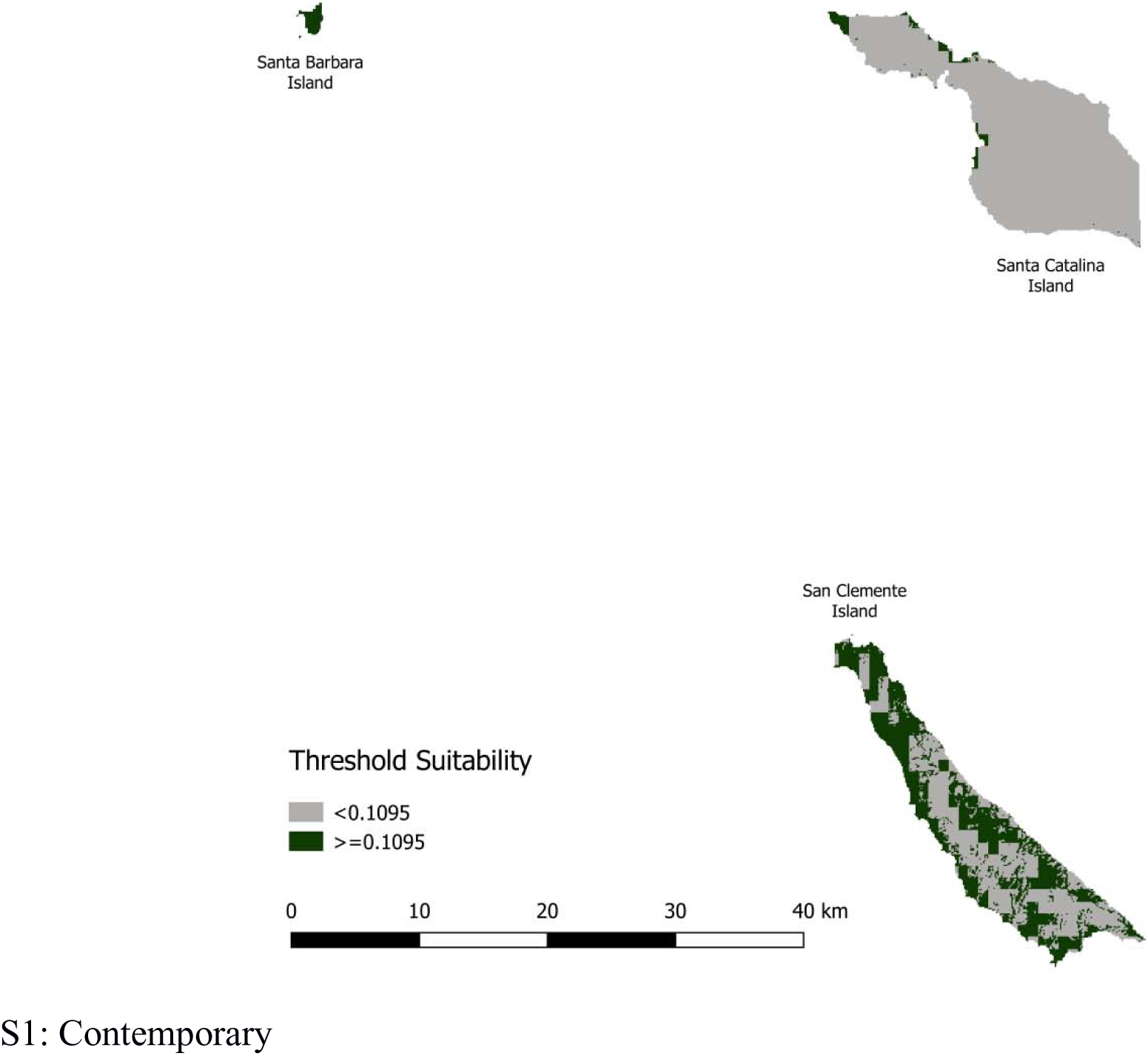
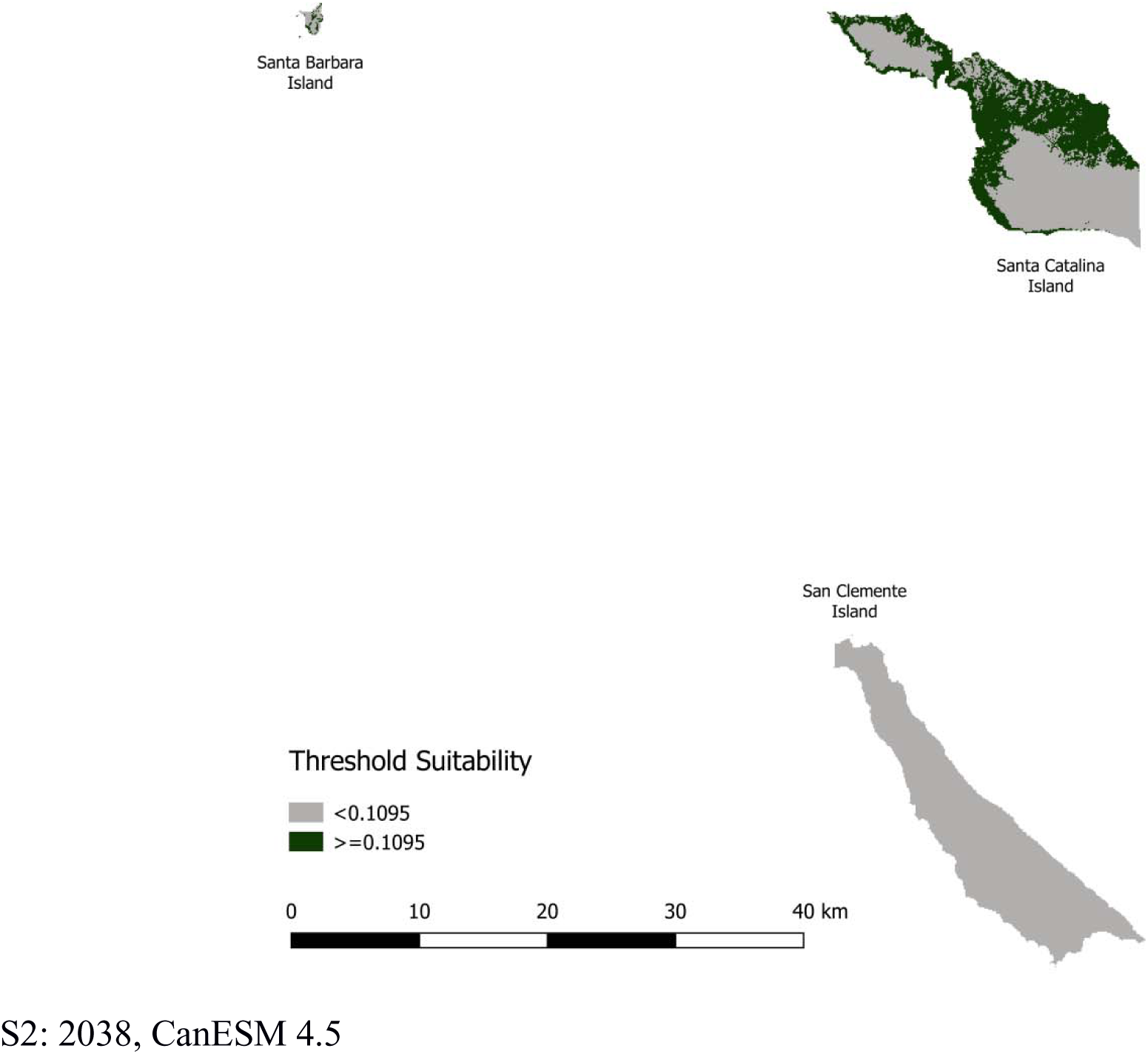
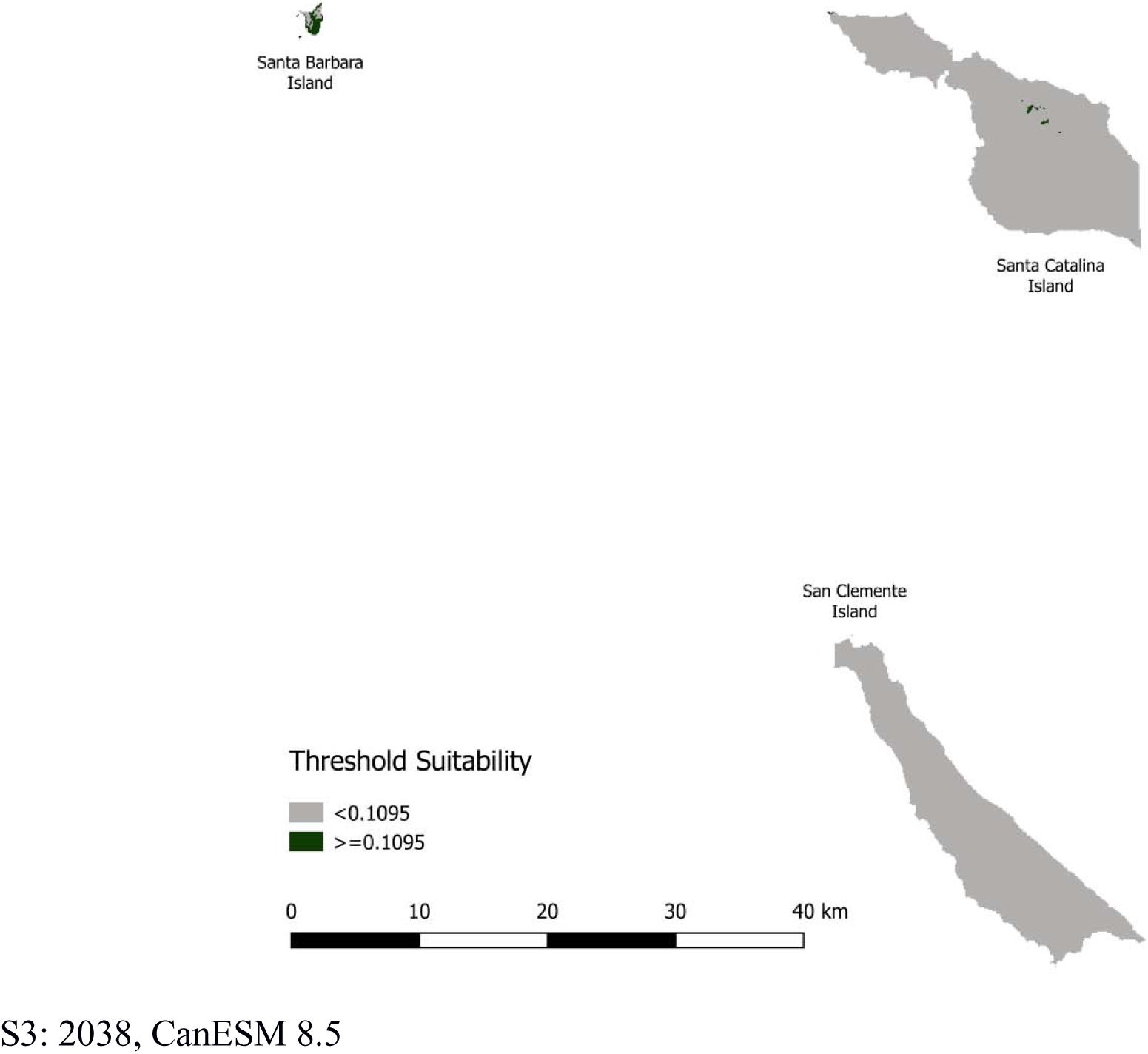
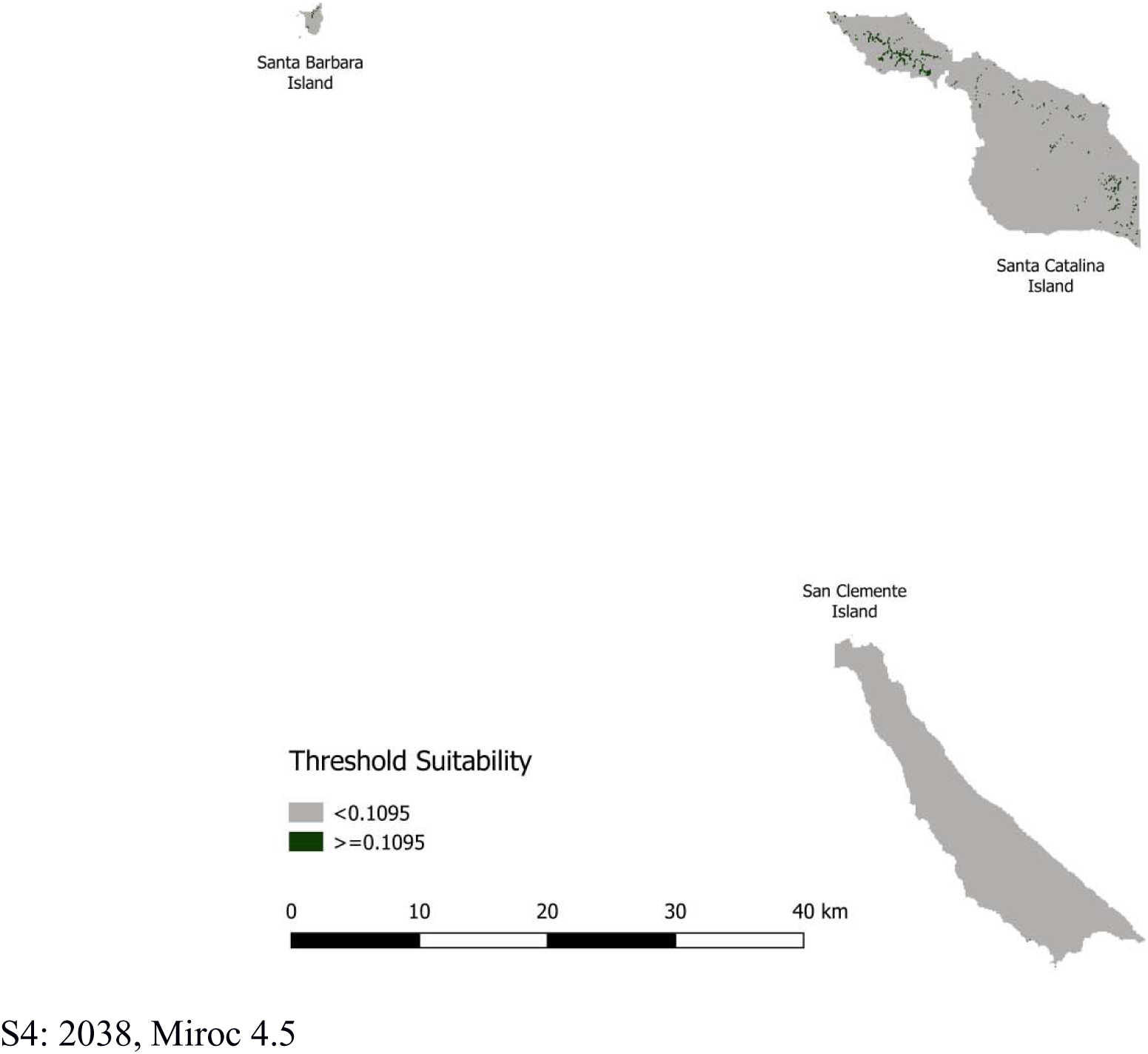
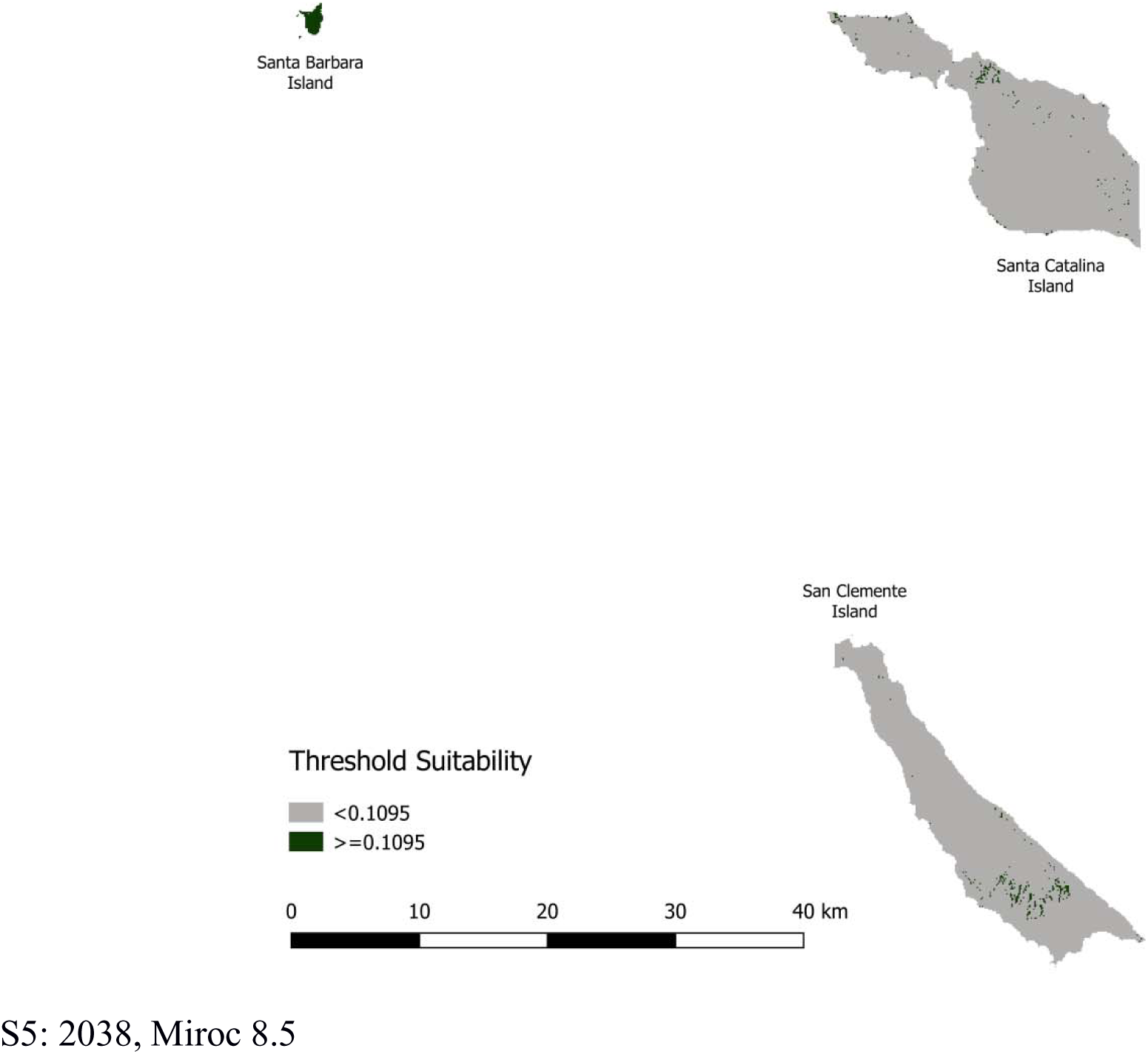
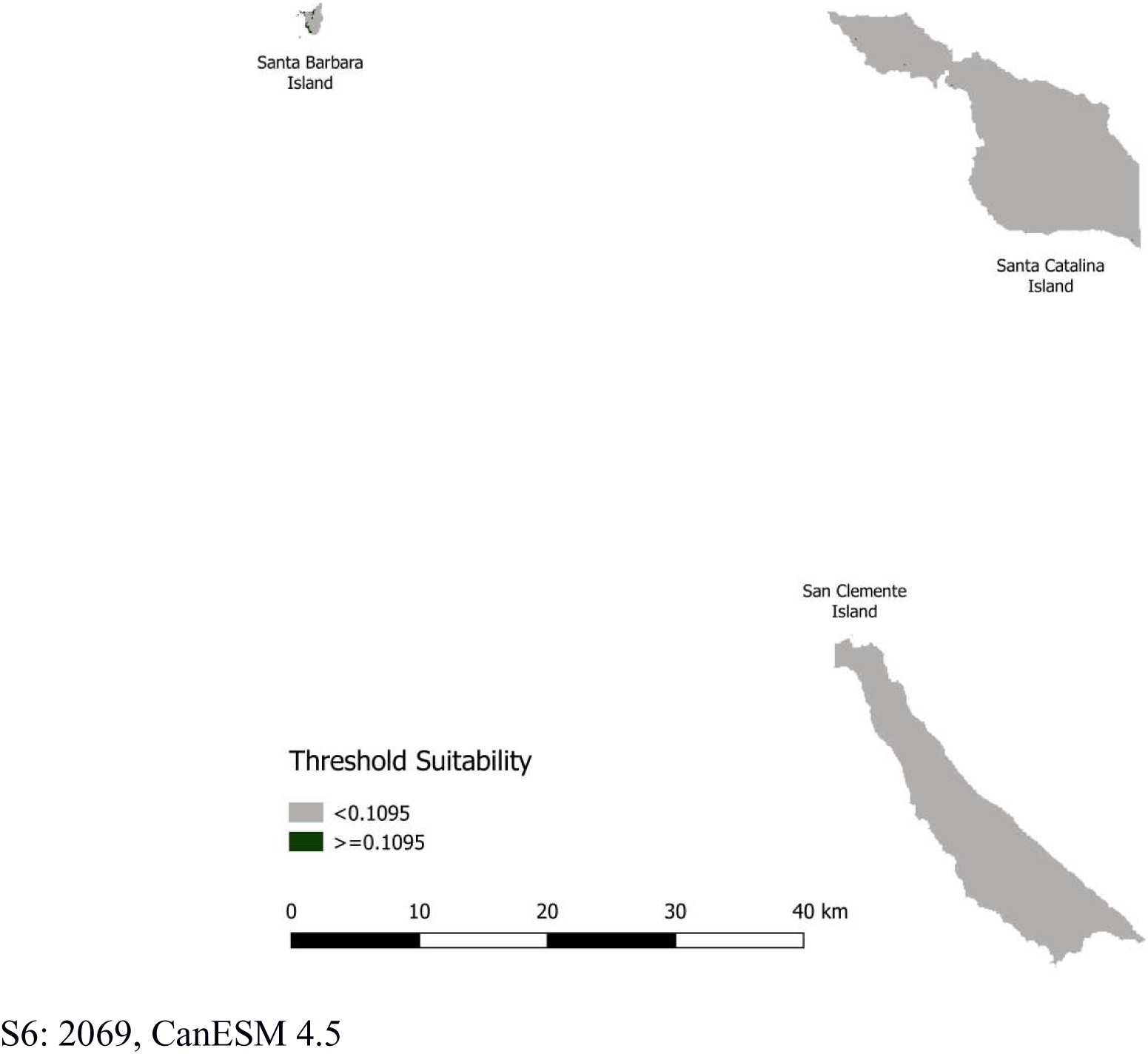
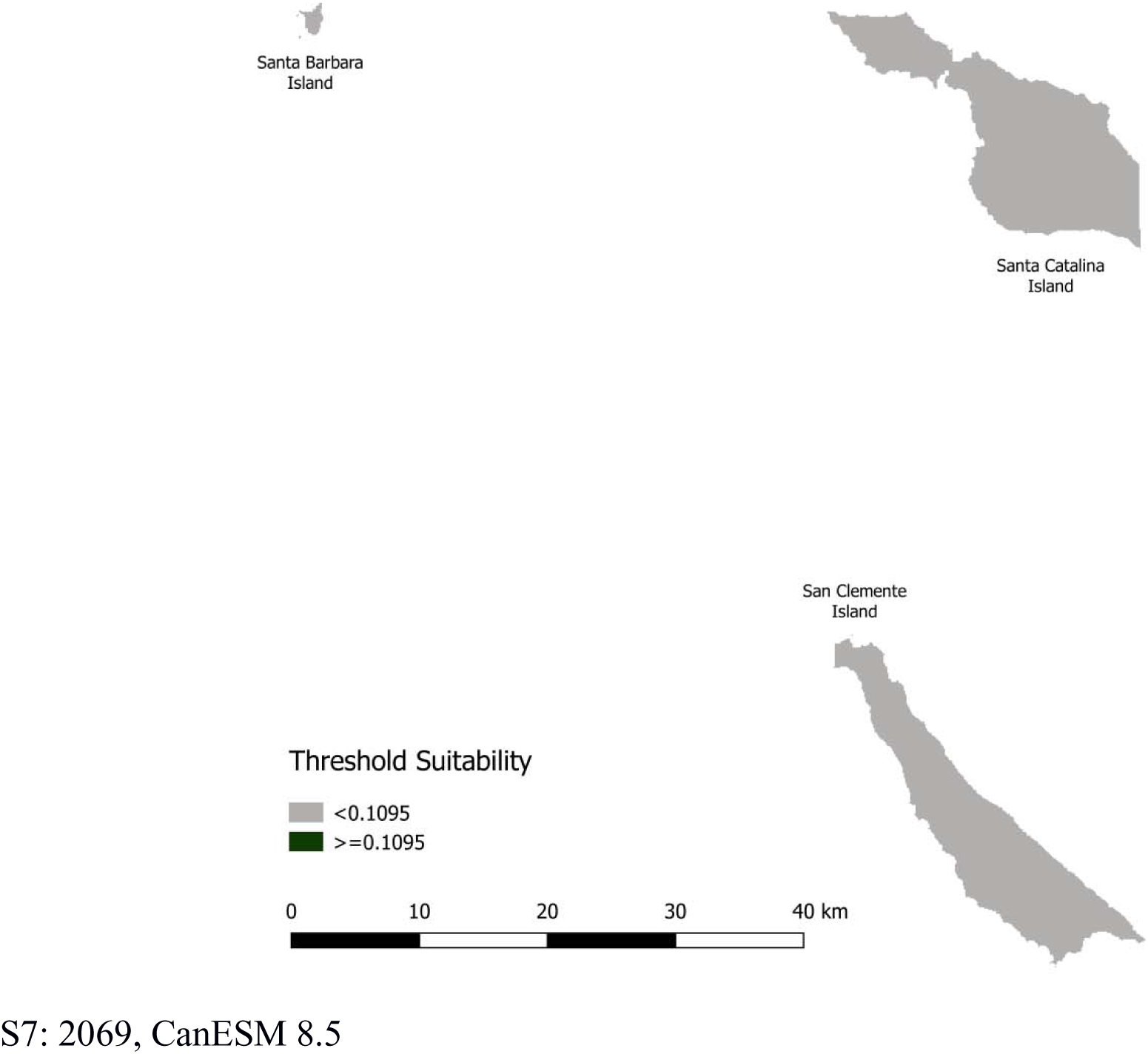
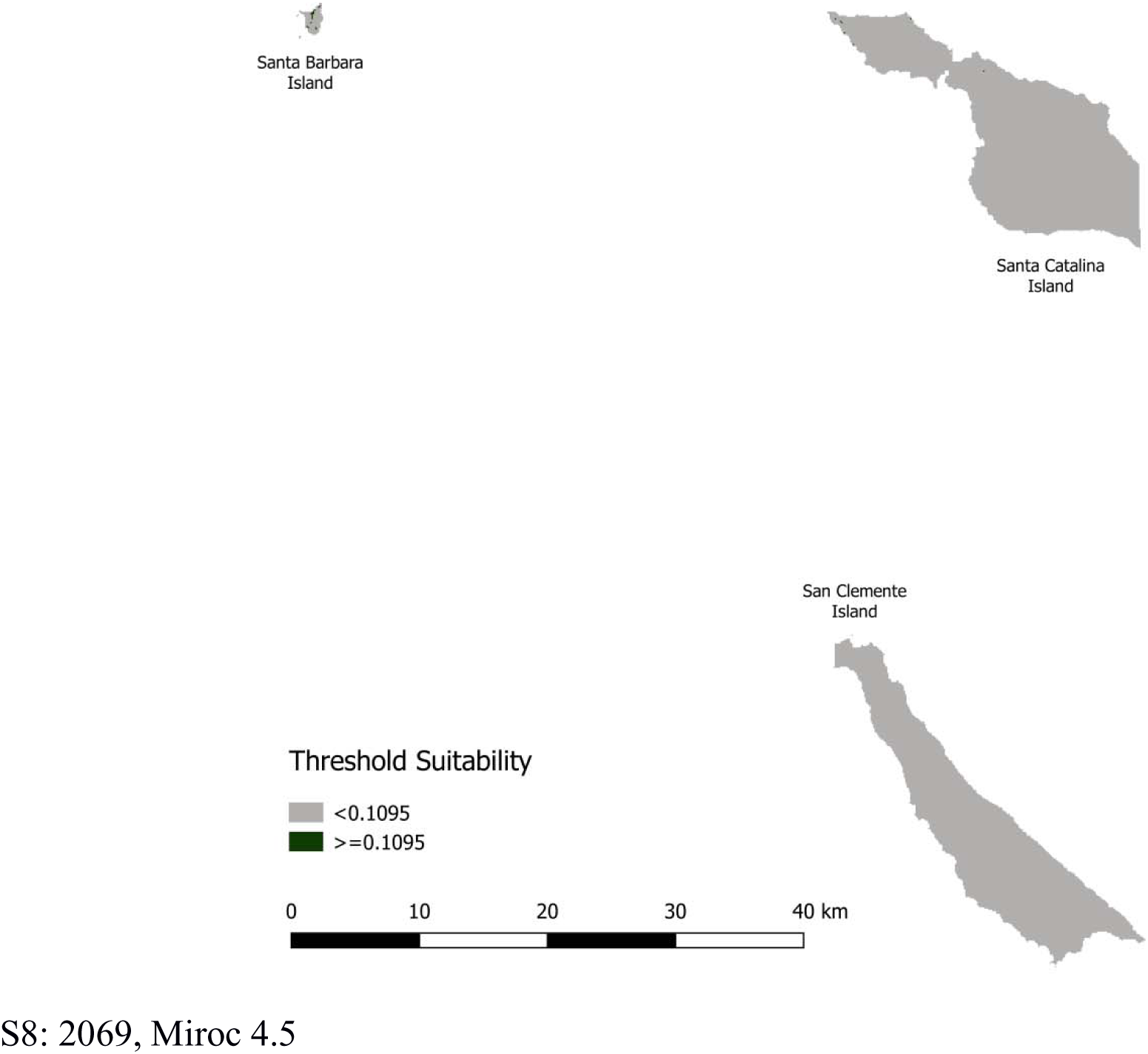
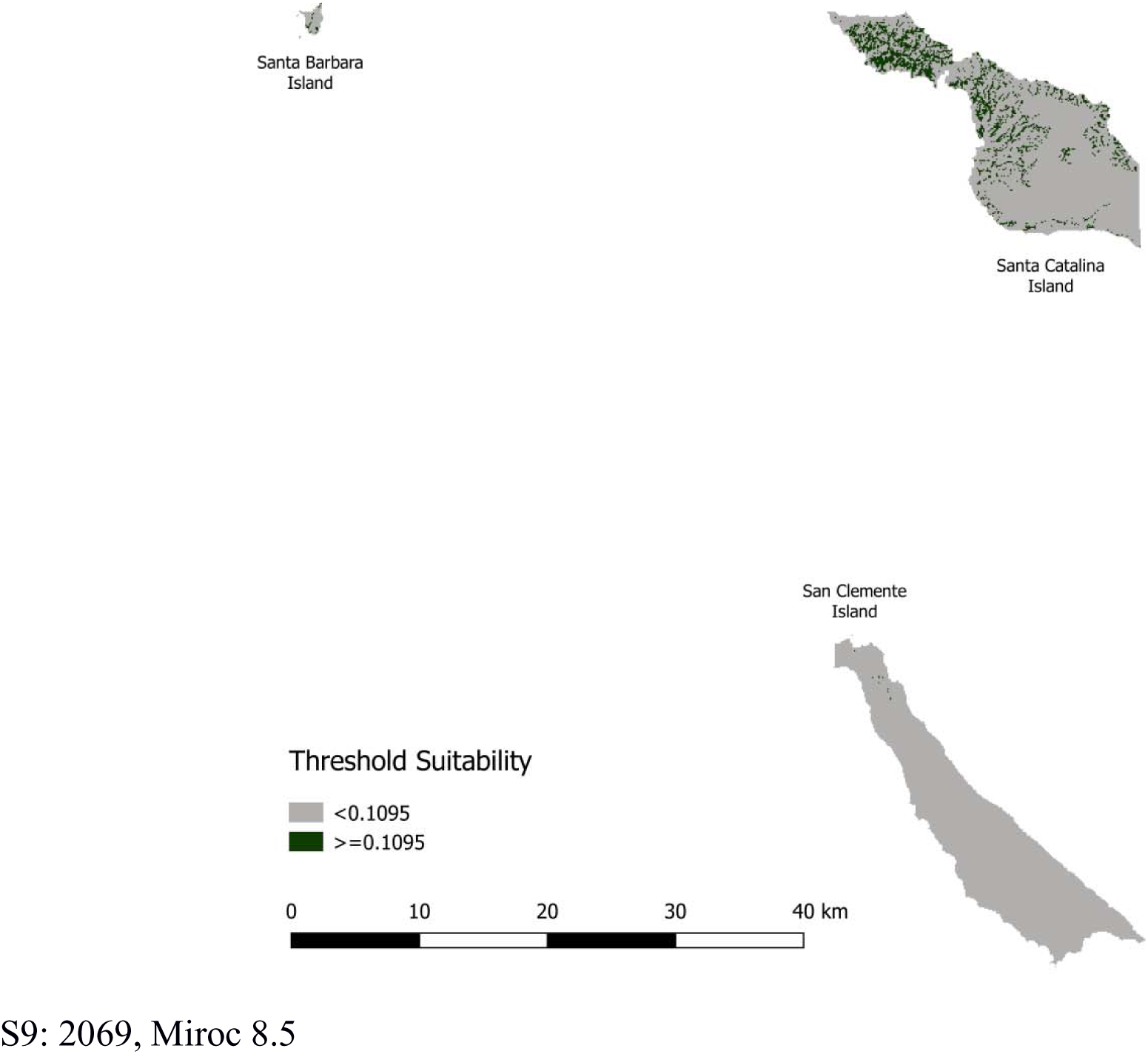
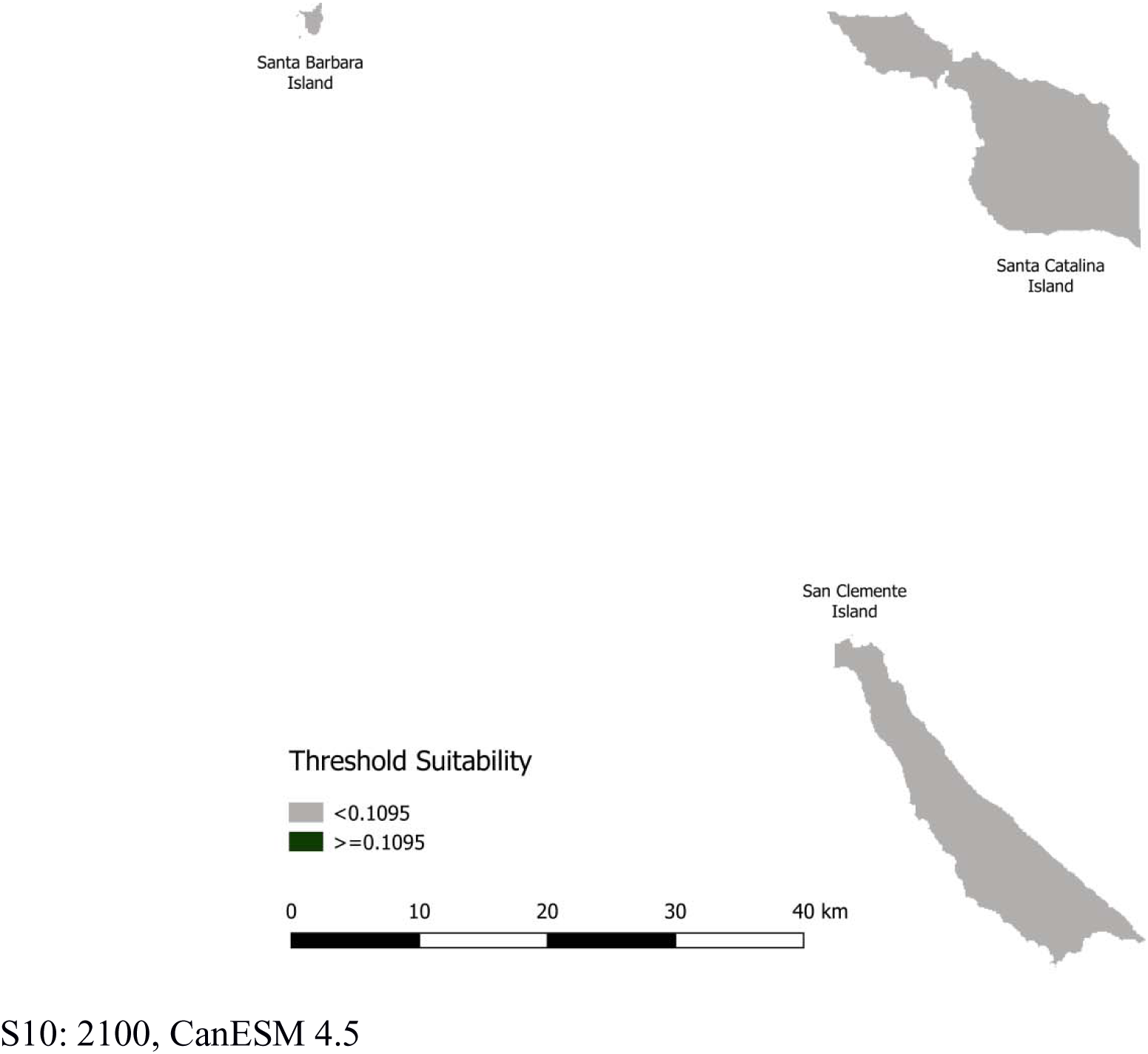
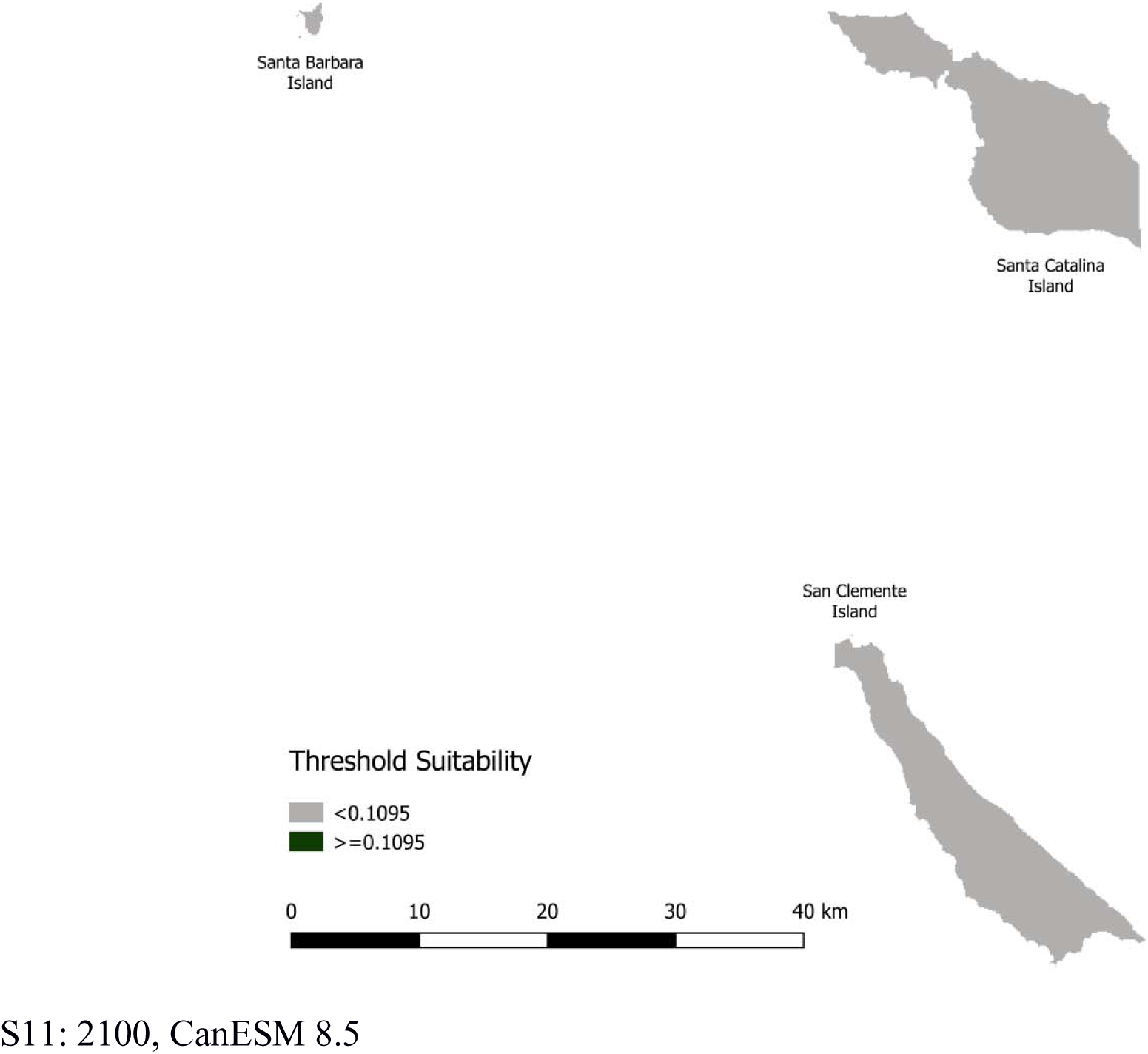
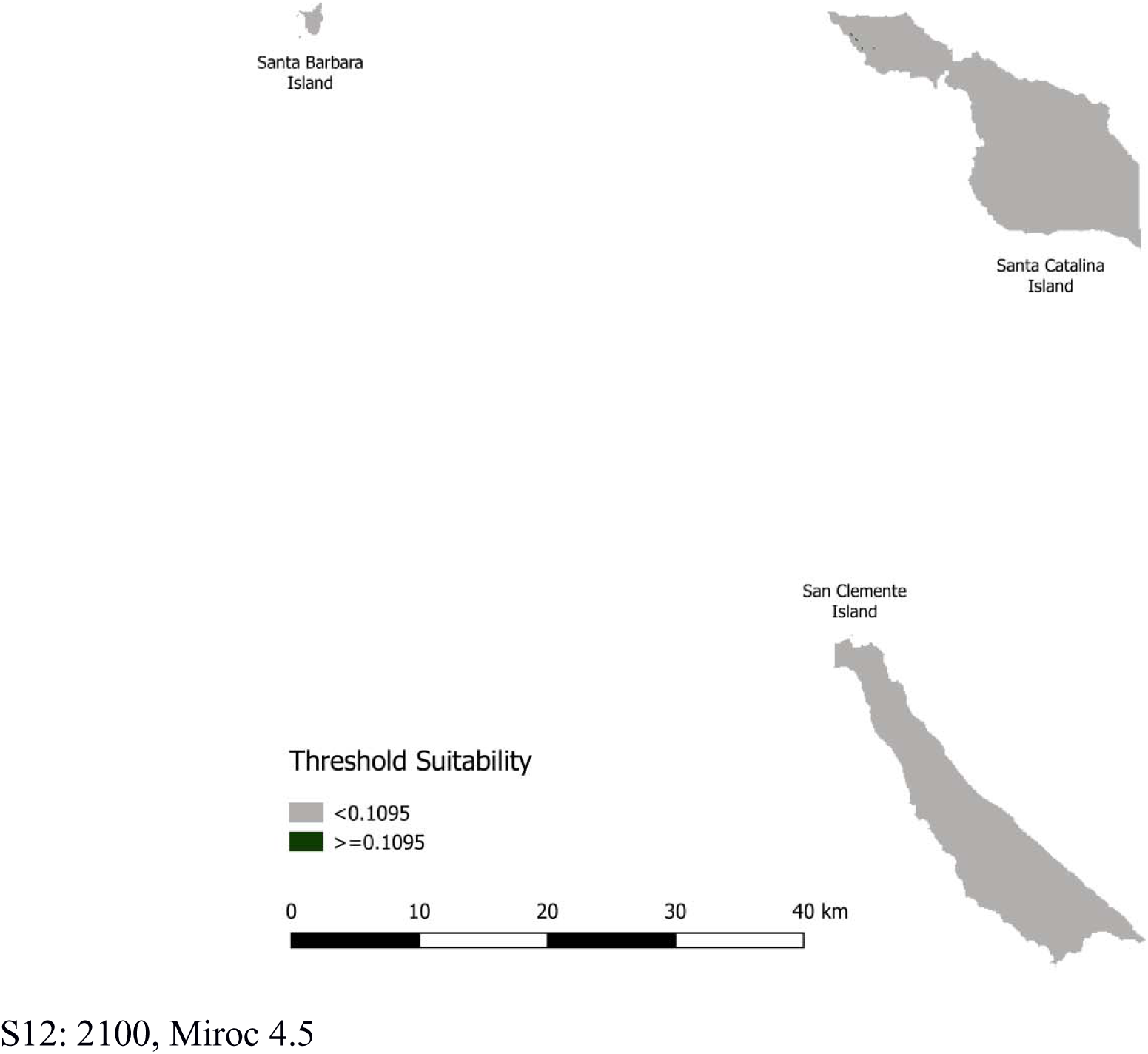
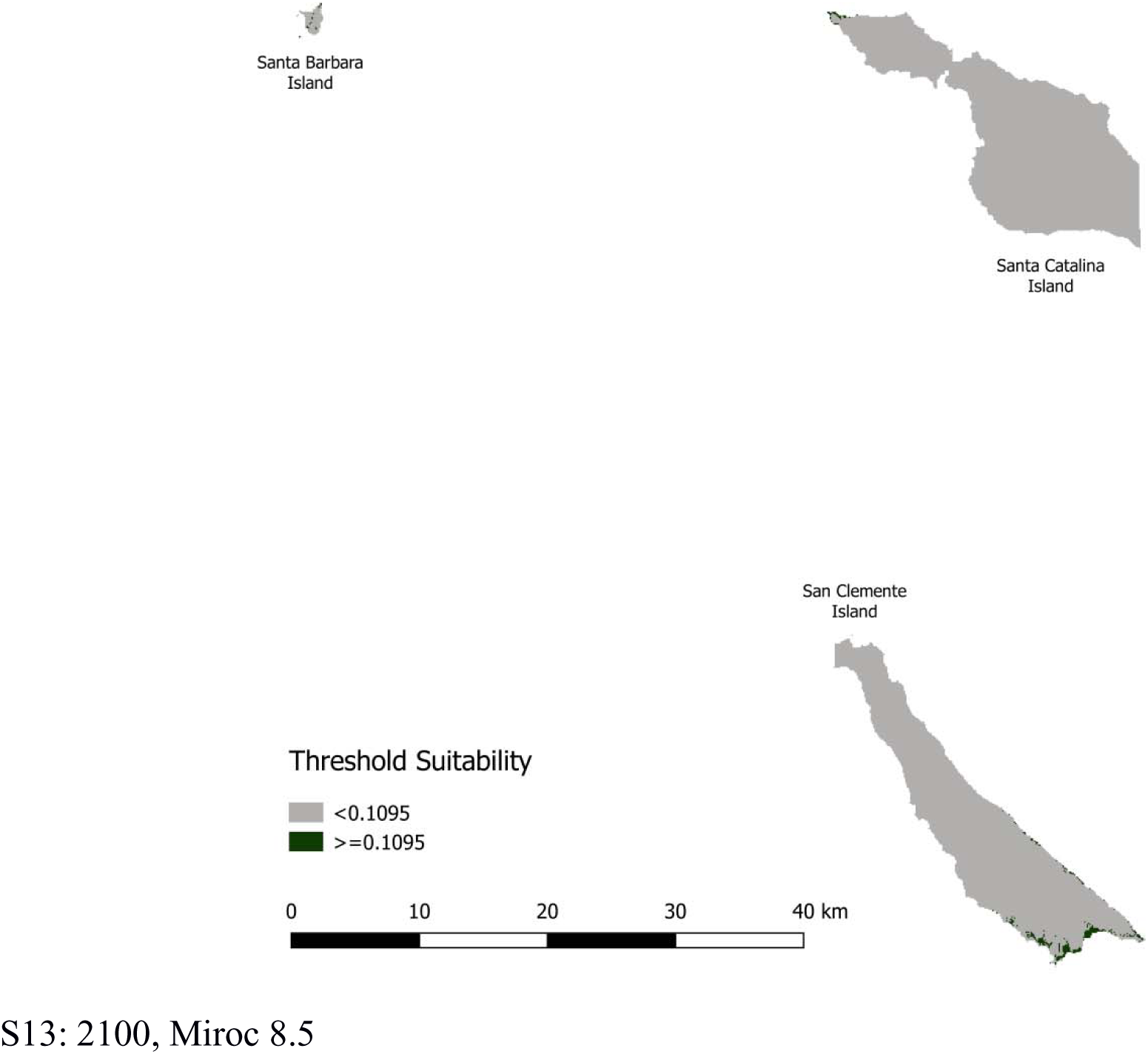
Habitat suitability models based on Threshold for all models in range. Models are identified by the endpoint, followed by the model name and RCP. Dark patches indicate suitable habitat based on the 0.1095 threshold value. Gray indicates unsuitably habitat below the threshold.

